# Euchromatin activity enhances segregation and compaction of heterochromatin in the cell nucleus

**DOI:** 10.1101/2022.02.22.481494

**Authors:** Achal Mahajan, Wen Yan, Alexandra Zidovska, David Saintillan, Michael J. Shelley

## Abstract

The large-scale organization of the genome inside the cell nucleus is critical for the cell’s function. Chromatin – the functional form of DNA in cells – serves as a substrate for active nuclear processes such as transcription, replication and DNA repair. Chromatin’s spatial organization directly affects its accessibility by ATP-powered enzymes, e.g., RNA polymerase II in the case of transcription. In differentiated cells, chromatin is spatially segregated into compartments – euchromatin and heterochromatin – the former being largely transcriptionally active and loosely packed, the latter containing mostly silent genes and densely compacted. The euchromatin/heterochromatin segregation is crucial for proper genomic function, yet the physical principles behind it are far from understood. Here, we model the nucleus as filled with hydrodynamically interacting active Zimm chains – chromosomes – and investigate how large heterochromatic regions form and segregate from euchromatin through their complex interactions. Each chromosome presents a block copolymer composed of heterochromatic blocks, capable of crosslinking that increases chromatin’s local compaction, and euchromatic blocks, subjected to stochastic force dipoles that capture the microscopic stresses exerted by nuclear ATPases. These active stresses lead to a dynamic self-organization of the genome, with its coherent motions driving the mixing of chromosome territories as well as large-scale heterochromatic segregation through crosslinking of distant genomic regions. We study the stresses and flows that arise in the nucleus during the heterochromatic segregation, and identify their signatures in Hi-C proximity maps. Our results reveal the fundamental role of active mechanical processes and hydrodynamic interactions in the kinetics of chromatin compartmentalization and in the emergent large-scale organization of the nucleus.

## I. INTRODUCTION

The genome stores information essential for life. This information exists in the form of the DNA molecule, which together with histone proteins, forms the chromatin fiber inside the cell nucleus [1, 2]. The chromatin fiber is intricately folded into a hierarchical threedimensional structure that is dynamical in nature [3, 4]. Its organization and dynamics directly affect vital processes such as transcription, replication and DNA repair [5–7], yet the physical principles underlying chromatin’s dynamical self-organization remain elusive.

In differentiated cells, the genome is further organized into euchromatin and heterochromatin compartments, comprised of predominantly transcriptionally active and inactive genes, respectively [4, 8]. The former is packed loosely, hence readily accessible to the transcription machinery, while the latter is densely packed [9, 10]. In most nuclei, heterochromatin forms throughout the nuclear interior and is enriched near the nuclear periphery, where it binds to the lamin filament network at the nuclear envelope [11, 12]. Notably, this organization is inverted in nuclei of photoreceptor cells of some nocturnal mammals, where chromatin–lamin interactions are missing [13, 14].

The biochemistry of heterochromatin formation is rooted in gene silencing associated with the methylation of histone tails (H3K9me3), which in turn enables the binding of heterochromatin protein 1 (HP1) [15–18]. Bound HP1 proteins then dimerize, bridging nearby chromatin fibers [19–22]. Remarkably, HP1 proteins have been found to undergo liquid-liquid phase separation from the nucleoplasm, forming liquid condensates surrounding the heterochromatin [23, 24]. In contrast to its biochemistry, physical principles behind the heterochromatin formation are far from understood.

Recent computational models have provided considerable insights into chromatin compartmentalization, by employing equilibrium simulations with chromatin represented as a block co-polymer, with euchromatin and heterochromatin segments having different respective affinities. Brownian dynamics and Monte Carlo simulations suggest that heterochromatin forms dense aggregates in the center of the nucleus in the absence of attractive interactions with the nuclear envelope [25, 26]. The addition of attractive interactions with the nuclear envelope (lamins) results in heterochromatin accumulation near the boundary [25, 27]. This is consistent with heterochromatin localization observed in conventional nuclei, confirming a key role of chromatin–lamin interactions in genome organization [14, 28]. These models considered equilibrium statistics, thus neglecting the role played by ATP-driven active processes in the kinetics of heterochromatin formation and its final morphology. Equilibrium polymer simulations, which account for activity using an effective temperature, suggest that local differences in chromatin activity may lead to genome compartmentalization [29, 30].

The importance of activity in nucleus-wide chromatin dynamics was first discovered in experiments by Zidovska *et al*. [31], who developed displacement correlation spectroscopy (DCS) to quantify chromatin positional dynamics simultaneously across the entire nucleus in live human cells by imaging histones H2B-GFP. These observations revealed that chromatin moves coherently within large regions of 3–5 *μ*m for several seconds. These correlated motions were found to be ATP-dependent, but independent of the cytoplasmic cytoskeleton [31]. Perturbation of major nuclear ATPases such as DNA polymerase, RNA polymerase II and topoisomerase II caused local displacements to increase and eliminated the large-scale coherence [31].

Motivated by these experiments, Bruinsma *et al*. [32] proposed a hydrodynamic theory of chromatin dynamics that coarse-grained the action of nuclear ATPases into two types of active events: vector events that describe force dipoles generated by nuclear enzymes such as polymerases and helicases, and scalar events corresponding to the local de/condensation of chromatin caused by chromatin remodelers. This mean-field continuum theory predicts that dipolar activity is responsible for large-scale coherent motions [31, 32]. An alternative hydrodynamics-free approach to modeling chromatin activity has been developed using a 3D-conformational space of the chromatin fiber, which emerges from a quasi-equilibrium energy landscape generated by Langevin dynamics at an effective temperature [33]. Another hydrodynamics-free model, informed by Hi-C data of human chromosomes, described chromatin as a heteropolymer and mimicked chromatin activity as isotropic white noise [34, 35]. Both models successfully recapitulated large-scale coherence of chromatin motions.

In recent work, Saintillan, Shelley and Zidovska [36] built upon the concept of vector events to develop a computational model of active chromatin hydrodynamics that accounts for the role of ATP-driven processes. In this model, a long flexible polymer was confined inside a spherical cavity, immersed in a viscous fluid (nucleoplasm) and subjected to stochastic force dipoles along the polymer. These force dipoles generated fluid flows in the surrounding nucleoplasm, which drove a local nematic alignment of the polymer chain and consequently of the dipoles as well [36]. Moreover, such dipolar forces were found to lead to spontaneous straightening of the polymer [37]. This positive feedback occurred only for extensile dipoles and caused large-scale coherent motions of the polymer by a mechanism similar to the generic bend instability of extensile active nematics [38]. These simulations assumed a homopolymer chain, thus no macroscopic self-organization was observed beyond the alignment induced by hydrodynamic interactions [36].

Here, we extend this computational approach to investigate the role of dipolar activity and nucleoplasmic flows in the spatial segregation of euchromatin/heterochromatin in differentiated cells. To do so, we perform large-scale coarse-grained simulations of a model nucleus containing multiple chromosomes immersed in a viscous nucleoplasm. The chromosomes are modeled as diblock copolymers composed of segments of active euchromatin subject to dipolar activity, alternating with segments of inactive heterochromatin subject to inter- and intra-chain crosslinking. Our numerical results reveal the role of activity-driven coherent motions and nucleoplasmic flows on the formation, spatial distribution and compaction of heterochromatin inside the nucleus. We present the coarse-grained chromatin model used in our simulations in Sec. II, and a detailed discussion of computational methods in Appendix A. Numerical results and their predictions for the spatial organization of euchromatin and heterochromatin are presented in Sec. III, where we show stress distributions and hydrodynamic flows due to chromatin activity as well as examine their consequences for the observed genomic contact probabilities. We discuss these results and their implications in detail in Sec. IV.

## II. COARSE-GRAINED CHROMATIN MODEL

We simulate a system of *M*_*c*_ confined polymers representing individual chromosomes inside the interphase nucleus (Fig. 1). The nuclear envelope, which encloses the chains, is modeled as a prolate spheroidal cavity with bounding surface *S* of eccentricity *e* and equivalent radius *R*_*s*_, and is filled with a viscous Newtonian nucleoplasm with viscosity *η*. Each chromosome is coarse-grained as a self-avoiding active Zimm bead-spring chain composed of *N*_*b*_ hydrodynamically interacting beads of radius *a*_*h*_, which are connected by finitely extensible elastic springs. In this coarse-grained description, a bead should be viewed as a mesoscopic chromatin sub-domain containing a large number of nucleosomes and associated proteins at a resolution of ∼ 50 kbp. As depicted in Fig. 1, each chromosome further consists of alternating blocks of heterochromatin (HC) and euchromatin (EC), with an HC fraction *α*_*c*_. Transcriptionally active euchromatin experiences microscopic active stresses generated by the ATP-powered action of nuclear enzymes such as RNA polymerase II. We model these stresses in terms of stochastic active force dipoles distributed along EC blocks and applied to the viscous fluid in which they drive an active flow [36]. On the other hand, the transcriptionally inactive HC blocks are characterized by the absence of active force dipoles and by the ability to form inter- and intra-chain crosslinks, which represent a macroscopic model for the crosslinking of the HC blocks by the heterochromatin protein 1 (HP1) [20, 26, 39].

**FIG. 1.**
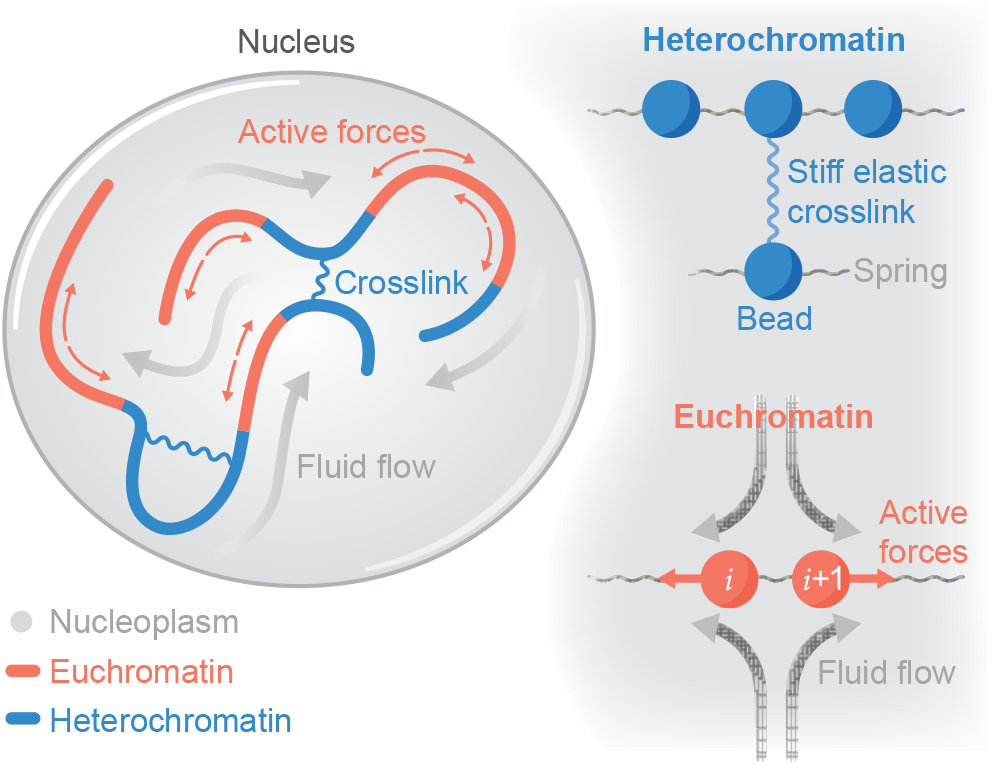
Model schematic: *M*_*c*_ bead-spring polymer chains representing individual chromosomes are confined inside a spheroidal nuclear envelope. Heterochromatic (HC) blocks can form inter- and intra-chromosomal crosslinks, whereas euchromatic (EC) blocks are decorated with active force dipoles that drive nucleoplasmic flows.

The position **x**_*i*_(*t*) of bead *i* = 1, …, *N* (where *N* = *N*_*b*_ *× M*_*c*_) is governed by the overdamped Langevin equation [40]

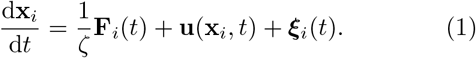

The first term on the RHS is the velocity contribution arising from the net deterministic force on bead *i*, to be specified below, where *ζ* = 6*πηa*_*h*_ is the corresponding friction coefficient. The second term captures advection of the bead by the viscous flow inside the nucleus, with contributions from activity as well as hydrodynamic interactions resulting from viscous drag on the beads. Finally, the last term captures Brownian displacements and is calculated to satisfy the fluctuation–dissipation theorem.

The net deterministic force **F**_*i*_ on bead *i* accounts for entropic spring tensions, crosslink forces as well as excluded volume interactions between neighboring beads:

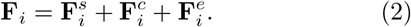

Entropic spring forces 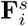 are modeled using the FENE spring law [41], and excluded volume forces 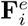 are captured using a soft repulsive potential; see Appendix A for details. Crosslink forces 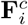 are used to model binding interactions between beads that belong to HC blocks. These crosslinks represent HP1 dimers bridging two heterochromatic fibers, their binding facilitated by the histone methylation H3K9 of heterochromatin [19–22]. These binding interactions, which can occur inter- or intra-chain, are also captured using FENE springs but with a stiffer spring constant. These links can form stochastically between pairs of heterochromatic beads that are within a cutoff radius *a*_*c*_ of one another and both are in an activated state, where activation occurs stochastically as a Poisson process with rate 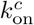. Each bead can form at most one crosslink, and crosslinks are permanent once formed.

A key aspect of our model is the inclusion of hydrodynamic flows induced by the activity of ATP-powered enzymes and from interactions between chain segments. Following our past work [36], microscopic active stresses are coarse-grained in the form of active dipolar forces applied to the fluid along euchromatic portions of the chromosomes and occur on the scale of one chain link. Specifically, active dipoles are switched on and off stochastically as a Poisson process with rates 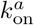 and 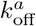, which set the probability 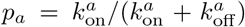 for a link to be active. When a euchromatic link *i* is in the on–state, two equal and opposite forces are applied to the viscous nucleoplasm at the positions of the end beads:

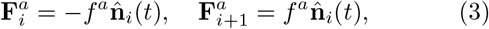

where 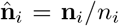 is the unit vector pointing from bead *i* to *i* + 1 obtained from the connector **n**_*i*_ = **x**_*i*+1_ **x**_*i*_ between the two beads. The pair of forces constitutes a dipole of magnitude *σ* = *f*^*a*^*n*_*i*_, which is extensile (*←→*) for *σ >* 0 and contractile (*→←*) for *σ <* 0. In simulations, we fix *f*^*a*^, meaning that the actual dipole strength fluctuates with the distance *n*_*i*_ between the beads. We define a characteristic dipole strength as *σ*_0_ = *f*^*a*^*𝓁*_*s*_, where *𝓁*_*s*_ is the equilibrium length of an isolated passive spring.

Both passive and active forces contribute to the nucleoplasmic flow field **u** and corresponding pressure *p*, which satisfy the Stokes momentum equation

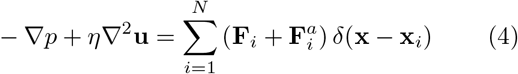

along with the incompressibility constraint ∇ · **u** = 0. In Eq. (4), active forces on the right-hand side are only included for beads that are in the active state. The velocity field is subject to the no-slip condition **u** = **0** on the surface of the nuclear envelope and is obtained using an accelerated algorithm based on the kernel-free fast-multipole method [42], allowing us to simulate large systems over long periods of time. In the following, we present results in dimensionless form based on characteristic thermal (equilibrium) scales. Further details of the computational methods and scalings are provided in Appendix A.

## III. RESULTS

In this section, we consider simulations in a spheroidal nucleus of equivalent radius *R*_*s*_ = 28, filled with *M*_*c*_ = 23 identical chains (or chromosomes) of 1305 beads for a total of *∼* 30k beads, with each bead representing *∼* 50 kbp of the human haploid genome. Each chromosomal chain contains four equally spaced linear HC blocks that in total occupy a fraction *α*_*c*_ *≈* 30% of each chromosome. Recall that crosslinks can form within and between HC blocks, but not with or within the EC blocks. Each chromosome is initially prepared as a confined random walk, then all chromosomes are distributed within the nucleus to establish distinct chromosomal territories. This system is then equilibrated under thermal fluctuations and excluded volume forces to establish initial data. Simulations were also performed on smaller systems, in which the chromosomes were initially prepared as fractal globules [44] instead of random walks (not shown), with no discernible differences found in the large-scale dynamics discussed below.

### A. Dynamics of heterochromatin segregation

Figure 2 shows the results of two long-time simulations proceeding from identical initial data (top row). In the first case, the EC blocks are entirely passive (*σ*_0_ = 0, middle row), and the dynamics largely results from crosslinks forming within and between HC blocks. In the second case, the EC blocks are active (*σ*_0_ *>* 0, bottom row), being stochastically populated with extensile dipoles; see videos of the dynamics in the Supplemental Material [43].

**FIG. 2.**
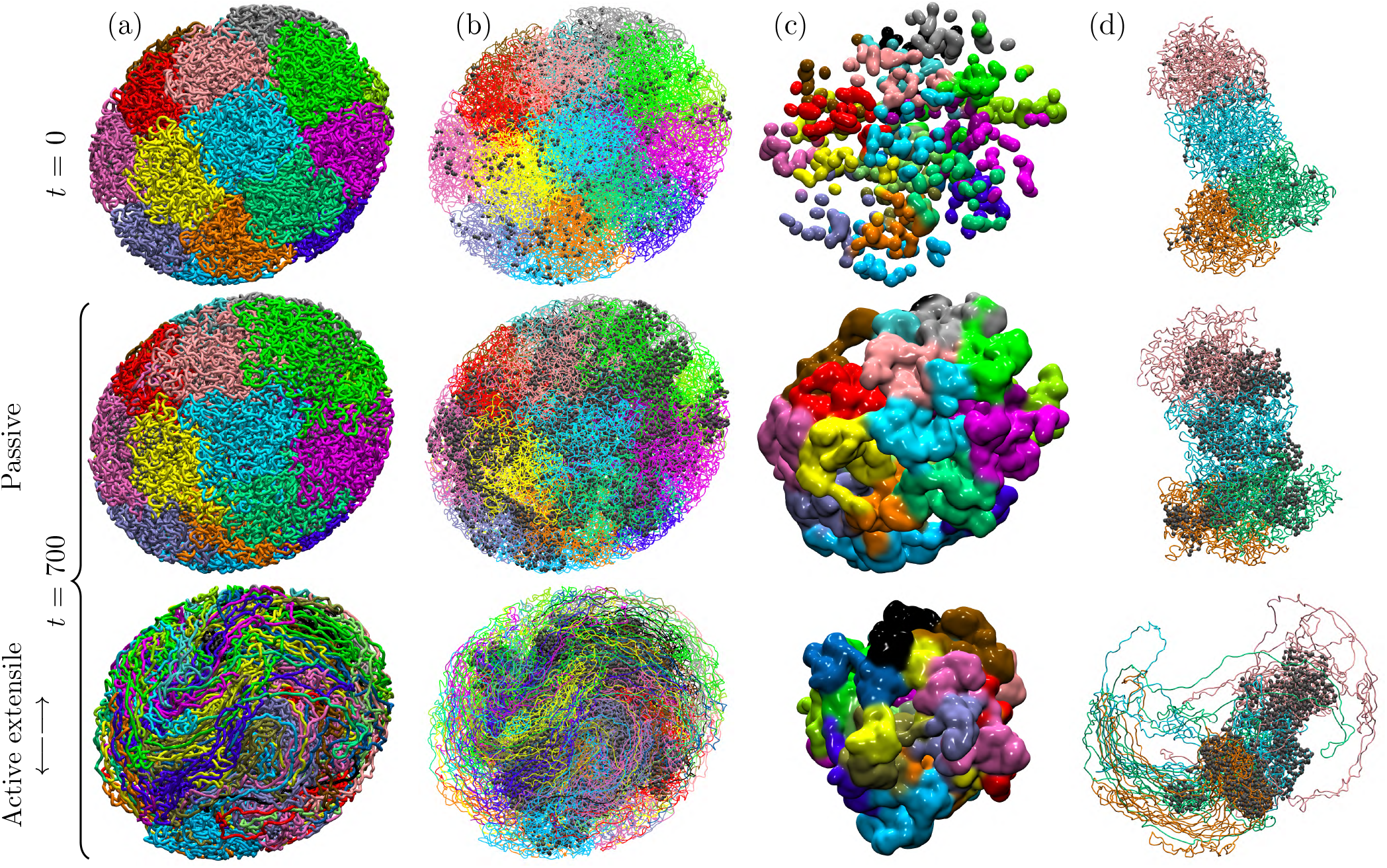
Long-time evolution of chromatin in simulations of two systems, one with passive euchromatin (*σ*_0_ = 0) and another with active euchromatin (extensile, *σ*_0_ = 30). Top row: initial system configuration at *t* = 0, which is identical in both simulations; second and third rows: system configurations at dimensionless time *t* = 700. The four columns show, from left to right: (a) the full chromatin system, with colors corresponding to distinct chromosomes; (b) the crosslinks within and between HC blocks, shown as grey beads and overlayed on top of the chains; (c) the location of HC regions (HCRs), defined as regions in space where the crosslink density exceeds 50% of its maximum measured value in the simulations, and colored by chain membership; and (d) a subset of four chromosomes and associated crosslinks. Also, see videos of the dynamics in the Supplemental Material [43].

In the passive case, the chromosomes roughly maintain their initial spatial territories (Fig. 2(a)), with only slight mixing, mainly due to thermal fluctuations, taking place near their boundaries. The stochastically nucleating crosslinks, being long-lived, have led progressively to crosslink-rich regions (Fig. 2(b)). These form a loose network that spans the entire system and alternates with crosslink-free euchromatin. However, the dynamics of crosslinking, together with any attendant change in the mechanics of the chromatin in response to thermal fluctuations and excluded volume forces, has not led to any large-scale rearrangements of chromosomes or HC blocks. We suspect that, being rapid in comparison to rearrangements by thermal effects, crosslinking may be rigidifying the system, leading to fewer opportunities for crosslinking between distant HC blocks. At any rate, heterochromatin is distributed across the nucleus with approximate uniformity, corresponding to the initial placement of the HC blocks; see Fig. 2(c).

The dynamics and distribution of heterochromatin is very different for a system that contains active euchromatin (Fig. 2, bottom row). There, extensile dipoles in EC blocks drive large-scale coherent flows that draw neighboring chain segments into alignment, while mixing chromosomal territories. As a consequence of these active flows, the morphology of crosslinked heterochromatic regions (HCRs, defined as the regions in space where the crosslink density exceeds 50% of its maximum measured value in the simulations) is also quite different: mixing indeed facilitates crosslink formation between distinct HC blocks, resulting in a more compact crosslinked network that progressively densifies near the center of the nucleus. These distinct morphologies are especially visible in Fig. 2(c), where we show the spatial boundaries of HCRs. It is important to note that while each HC block represents a part of one chromosome, an HCR can contain HC blocks from several different chromosomes. We find that HCRs have a much more compact and dense structure in the active case, with most of the active euchromatin expelled from the nuclear center towards the nuclear envelope. Snapshots of four individual chromosomes are also shown in Fig. 2(d) and further highlight the role of active nucleoplasmic flows that promote the compaction of heterochromatin while opening up and unfolding active euchromatic chain segments.

We further quantify the spatiotemporal evolution of the system in Fig. 3. The migration of heterochromatin towards the nucleus center in the active case is evident in Fig. 3(a,left), showing the time evolution of the standard deviation 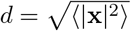 of the polymer mass distribution normalized by the equivalent nuclear radius *R*_*s*_ = 28, for both EC and HC blocks. While the spatial distribution of euchromatin shows only a minor shift towards the nuclear boundary, heterochromatin is found to concentrate near the center of the nucleus in the active extensile case, as evidenced by the decay of *d*_hc_ with time and corresponding growth of the migration offset defined as Δ = *d*_ec_ *− d*_hc_. This inward migration is also accompanied by an increase in chromatin number density inside heterochromatic regions as shown in Fig. 3(a,right), where we define *ρ*_hcr_ = *N*_hcr_*/V*_hcr_ as the ratio of the number of beads of any type contained in the HCRs identified in Fig. 2(c) over the corresponding volume. In the absence of activity (*σ*_0_ = 0), the number density remains close to the average number density of the system, whereas it increases significantly when extensile dipoles are applied along EC blocks (*σ*_0_ *>* 0). As shown by the insets in Fig. 3(a,right), this density increase is due to both an increase in *N*_hcr_ and a decrease in *V*_hcr_, suggesting that activity not only drives a spatial compaction of HCRs over time, but also causes these compacting regions to recruit and trap more EC block fragments during compaction. Spatial maps of the heterochromatic density in a plane cross-section are shown in Fig. 3(b) and confirm these findings: crosslinked regions are relatively sparse and evenly distributed across the nucleus in the passive case, but concentrate into a dense and compact structure near the center of the nucleus with extensile activity. The case of contractile activity (not shown) is qualitatively similar to the passive case, with no discernible segregation of HC and EC blocks.

**FIG. 3.**
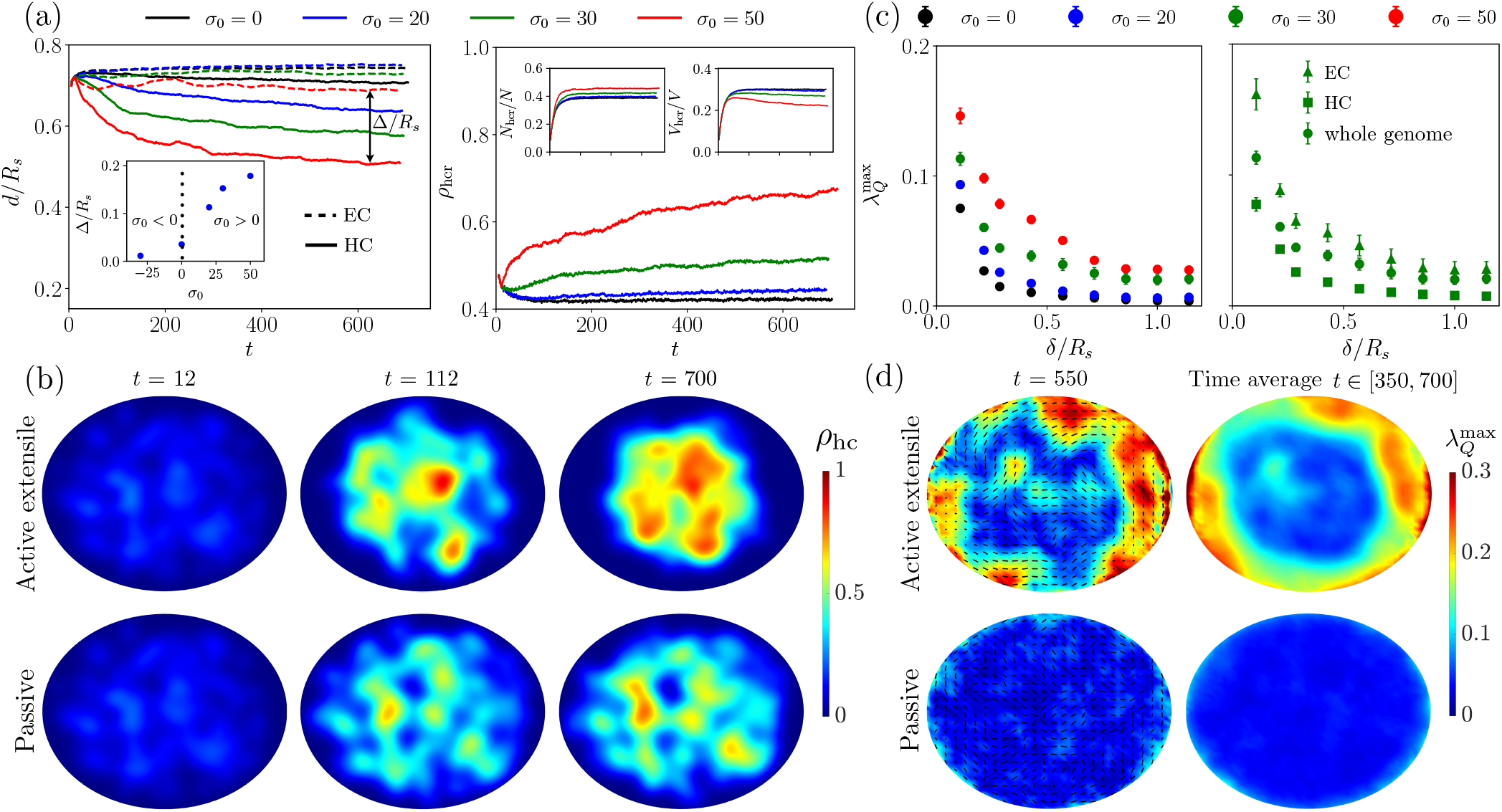
(a) Left: Standard deviation 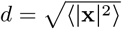 of the polymer mass distribution normalized by the equivalent nucleus radius *R*_*s*_ for EC and HC blocks as a function of time, where HC blocks are seen to migrate towards the center of the system. The inset shows the migration offset, defined as Δ = *d*_ec_ − *d*_hc_, at steady state as a function of activity *σ*_0_. Right: Mean density *ρ*_hcr_ of HCRs (see Fig. 2(c)). Insets show the fraction of the number of beads contained in HCRs, as well as the fraction of the total volume that they occupy. (b) For the simulations in Fig. 2, time snapshots of the heterochromatin density field in a plane containing the major axis of the nucleus, for the active euchromatin case (*σ*_0_ = 30, top row), and the passive euchromatin case (*σ*_0_ = 0, bottom row). (c) Left: nematic order parameter 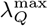, averaged over spheres of radius *δ* at late times, for various levels of activity *σ*_0_. Right: nematic order parameter for EC blocks, HC blocks, and for the whole genome, in simulations with *σ*_0_ = 30. (d) For the same simulations as shown in Fig. 2, the structure of nematic alignment for *δ* = 5 in a plane containing the major axis of the nucleus, for the active euchromatin case (*σ*_0_ = 30, top row) and the passive euchromatin case (*σ*_0_ = 0, bottom row). The colormap is of the scalar nematic order parameter, while black segments depict the nematic director projected onto the viewing plane. The left column shows a snapshot at *t* = 550, whereas the right column shows a time average for *t ∈* [350, 700].

As observed in the snapshots of Fig. 2, euchromatic extensile activity not only facilitates the segregation and compaction of crosslinked heterochromatin, but it also induces its own unfolding and alignment as first predicted in our past work [36, 37]. We quantify this alignment by calculating the tensorial nematic order parameter 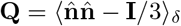, where 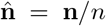 is the unit director between successive beads inside the chains, and the average is performed over spherical domains of radius *δ*. Its largest eigenvalue 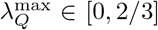 defines the scalar nematic order parameter and is a measure of the degree of euchromatin alignment on the length scale *δ*. Figure 3(c,left) shows 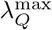 as a function of *δ* for different levels of activity. In passive simulations (*σ*_0_ = 0), nematic alignment is negligible except on short length scales (*δ/R*_*s*_ ≲ 0.2), where it is induced by steric interactions between neighboring chain segments. In the presence of extensile activity, the nematic order significantly increases on all scales as active force dipoles within EC blocks induce local flows that draw nearby chain segments into alignment. This emergent alignment increases the spatial coherence of the active dipolar flows via a positive feedback loop, generating nematic order on length scales that greatly exceed the scale of one dipole *∼ O*(1). The precise mechanism for these coherent flows was discussed in our past work [36] and is similar to the generic instability occurring in various other active nematic systems [45–47]. As shown in Fig. 3(c,right), nematic alignment primarily occurs among EC blocks, where dipolar activity takes place, as these sections of the chromatin are not crosslinked and therefore relatively free to reorganize in response to hydrodynamic flows. Alignment inside HC blocks is much weaker and comparable to the passive case, as the internal structure of heterochromatin is strongly constrained by the presence of crosslinks. These observations are amplified in Fig. 3(d), which shows spatial maps of the scalar order parameter and dominant alignment direction in a plane across the system. Nematic alignment is strong in euchromatic regions on the nuclear periphery and tends to conform to the system boundary. These features remain present in the time-averaged nematic order parameter map, where the average was performed over *t ∈* [350, 700]. The strong heterogeneity of the average map on this timescale suggests long-lived internal dynamics in the system, despite its very dynamic nature on short timescales.

In summary, our results reveal the central role of ATP-powered activity, taking place along the EC blocks of each chromosome, in determining the density, structure and positioning of heterochromatic regions (HCRs) inside the nucleus. We find that activity enhances the HCR compaction as well as the trapping of euchromatic segments within HCRs, relative to the passive case. Moreover, activity creates a large-scale nematic alignment outside of HCRs, where EC blocks are largely unconstrained and free to align with themselves and with boundaries of the nucleus as well as HCRs. The mechanistic origin of this complex organization lies in the emergent stress fields – both active and passive – and attendant nucleoplasmic flows, which we now analyze.

### B. Active stresses and hydrodynamic flows

The dynamics of heterochromatin region formation and euchromatin nematic ordering are tightly linked to the hydrodynamic flows driven by active stresses and the resulting elastic stresses they generate inside the chains. We first analyze the internal stress distributions in Fig. 4(a), showing maps of the radial (Σ_*rr*_) and shear (Σ_*rθ*_) components of the active and tensile stress tensors in a plane across the nucleus, where (*r, θ*) are polar coordinates in that plane. The distribution of active dipoles along EC blocks results in an effective active stress [48, 49] that can be defined, based on the Irving– Kirkwood formula [50], as

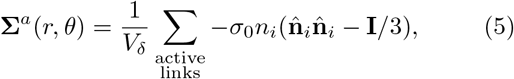

where the sum is over all active links inside the local averaging volume *V*_*δ*_, taken to be a sphere of radius *δ* = 5. Similarly, the tensile stress is calculated as a local average over the FENE springs comprising the chromatin and crosslinks as

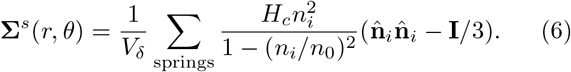

**FIG. 4.**
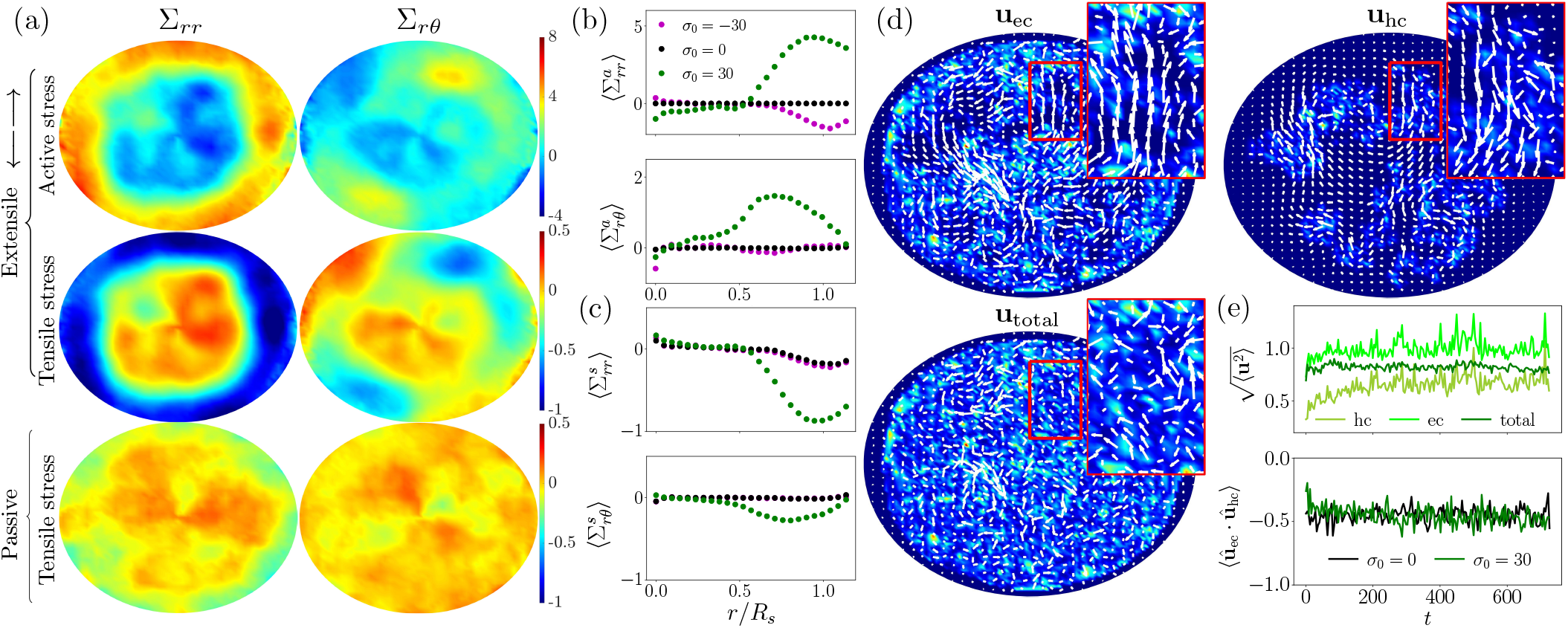
Analysis of the stress and flow fields: (a) Stress field exerted by dipolar active forces (first row) and internal tensile forces (second and third rows) in a plane containing the major axis of the nucleus, for the active euchromatin case (extensile, *σ*_0_ = 30, top two rows) and the passive euchromatin case (*σ*_0_ = 0, bottom row). The two columns show the radial (σ_*rr*_) and shear (Σ_*rθ*_) components of the stress averaged over *t ∈* [350, 700], where (*r, θ*) are polar coordinates in the plane. (b,c) Radial variation of active (b) and tensile (c) stress components averaged over the azimuthal direction for extensile (*σ*_0_ = 30), passive (*σ*_0_ = 0) and contractile (*σ*_0_ = *−* 30) systems. (d) Snapshots of the disturbance flows induced by deterministic forces along EC blocks (**u**_ec_) and HC blocks (**u**_hc_), as well as total disturbance flow (**u**_total_ = **u**_ec_ + **u**_hc_), for an active extensile system with *σ*_0_ = 30. The colormap shows the magnitude of the force distribution, and the vector plot shows the projections of the velocity field in a plane containing the major axis of the nucleus. Insets show zoomed-in regions where the flows induced by euchromatin and heterochromatin forces clearly oppose one another. (e) Top: Temporal evolution of the root mean square velocity 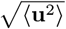, showing the stronger flow induced by the euchromatin due to the presence of active dipole forces. Bottom: Time evolution of the correlation between the directions of the euchromatin and heterochromatin velocity fields, defined as ⟨**û**_ec_ · **û**_hc_⟩ where **û** = **u**/|**u**|, showing a strong negative correlation. These panels use the data from the simulations in Fig. 2. Also see videos of the flow fields in the Supplemental Material [43].

In the active extensile case (*σ*_0_ *>* 0), active stresses are dominant in the nuclear periphery, where euchromatin is primarily located and organized, and their distribution closely follows that of the nematic tensor in Fig. 2(d). Indeed, in the mean-field limit, the active stress of Eq. (5) can be approximated as **Σ**^*a*^ *≈ − ρ*_*a*_*σ*_0_ ⟨*n*_*a*_⟩ **Q** where *ρ*_*a*_ is the local number density of active links and ⟨ *n*_*a*_ *⟩∼ O*(1) is the mean length of active links, which is activity dependent. In simulations with *σ*_0_ = 30, we measure ⟨*n*_*a*_⟩ ≈ 1.3. As shown by Fig. 4(a), a strong positive radial stress 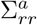 (similar to a negative active pressure) exists near the boundary and is consistent with the nematic alignment of the active extensile euchromatin chain segments along the boundary as found in Fig. 2(d). An active shear stress 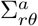 is also observed near the boundary, albeit less intense and not uniformly distributed, with both 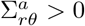 and *<* 0 over distinct euchromatic regions. As is evident in Fig. 4(a), the response of the elastic chromatin chain network to dipolar activity involves an oppositely signed tensile stress **Σ**^*s*^ with radial and shear components that closely mirror the active stress distribution. This indicates that the fluid flows driven by active stresses tend to stretch and shear the chromatin network resulting in the observed tensile stress. In the passive system, there are no active stresses and the tensile stress distribution shows no clear structure except very close to the system boundary, where weak alignment occurs due to steric effects. These various findings are summarized in Fig. 4(b,c) showing the radial dependence of the various stress contributions in extensile, passive, and contractile systems. In all cases, average stresses are quite weak inside heterochromatin (*r/R*_*s*_ *<* 0.5) but display a peak near *r/R*_*s*_ *≈* 0.7 *−* 0.9, where active stresses in extensile systems dominate the tensile stresses they induce inside the chain. With contractile activity, active stresses are of the opposite sign and are much weaker in magnitude due to the lack of nematic ordering in that case, other than very close to the nuclear envelope.

Figure 4(d) shows typical snapshots of the nucleoplasmic flow fields in the active extensile case; see videos of these flow fields in the Supplemental Material [43]. To analyze the distinct contributions of EC and HC blocks, we display separately the two flow fields induced by each type of chromatin as well as their sum, which is the net flow experienced by the system. The nucleoplasmic flow induced by euchromatin, which is distributed throughout the system, is found to be the strongest as a result of the active stresses present along these chain segments and is characterized by large-scale jets and recirculating flows. On the other hand, heterochromatin is primarily localized near the center of the system, and the disturbance flows it exerts are strongest there, where they are found to oppose the active flow driven by euchromatin. This is especially visible in the insets in Fig. 4(d), and is consistent with the discussion of stresses above: active flows tend to stretch and deform the heterochromatin network, which responds by developing tensile forces that oppose the flow. The net nucleoplasmic flow in the system, shown in the third panel of Fig. 4(d), is the sum of the two and indeed appears weaker and less coherent than the flow induced by euchromatin alone. This damping of active flows is indicative of hydrodynamic screening by the passive heterochromatic regions, which act as a porous elastic network. These findings are confirmed in Fig. 4(e), where we observe that the root mean square velocity of the total flow falls between those of the contributions from euchromatin and heterochromatin, and that the directions of these two velocity contributions tend to oppose one another: ⟨**û**_ec_ · **û**_hc_⟩ where **û** = **u**/|**u**|. Interestingly, the same negative correlation exists in the passive case, even though the motion in that case is purely thermal at steady state, and thus spatially uncorrelated and much weaker.

In summary, we find that the chromatin segregation into dense HCRs surrounded by nematically aligned euchromatin occurs concomitantly with the development of stress fields inside the nucleus. These stresses are strongest outside of HCRs and involve opposing contributions from euchromatic activity and internal tensile forces. The resulting fluid flows are coherent on large length scales and show evidence of hydrodynamic screening by the crosslinked HCR networks. These flows, together with crosslinking interactions, result in dynamic, yet highly structured chromatin conformations, whose impact on the genome organization we now examine.

### C. Hi-C proximity maps

The genome’s physical interactions can be quantified across different length scales by assessing the proximity probabilities between different genomic loci using chromosome conformation capture techniques such as Hi-C [51, 52]. We examine such interactions between chromosomes 1 and 4 in our model by measuring the proximity or near-contact of genomic loci along the two chromosomes. The resulting heat map (or Hi-C map) visualizing the contacts among genomic loci is shown in Fig. 5, with color ranging from red to white indicating high to low proximity, respectively (see Appendix B for details of the method used to calculate Hi-C maps). Figures 5(a,b) below the diagonal show snapshots at a late time for a passive (*σ*_0_ = 0) and active extensile simulation (*σ*_0_ = 30), respectively, whereas above the diagonal they show time averages over *t ∈* [350, 700] for the same simulations. The blue colored segments along the axes highlight the location of the HC blocks, while the green colored dots along the diagonal label portions of the genome that are spatially located inside the HCRs identified in Fig. 2(c). The relative size of these green high-interaction regions exceeds that of the HC blocks due to chain connectivity, which ensures that proximal genomic segments become trapped inside the HCRs as the permanent crosslinks form and compact the chain locally. This is especially true in the active extensile case (Fig. 5(b)) and is consistent with Fig. 3(a), which showed that the fraction of beads contained in HCRs increases with activity.

**FIG. 5.**
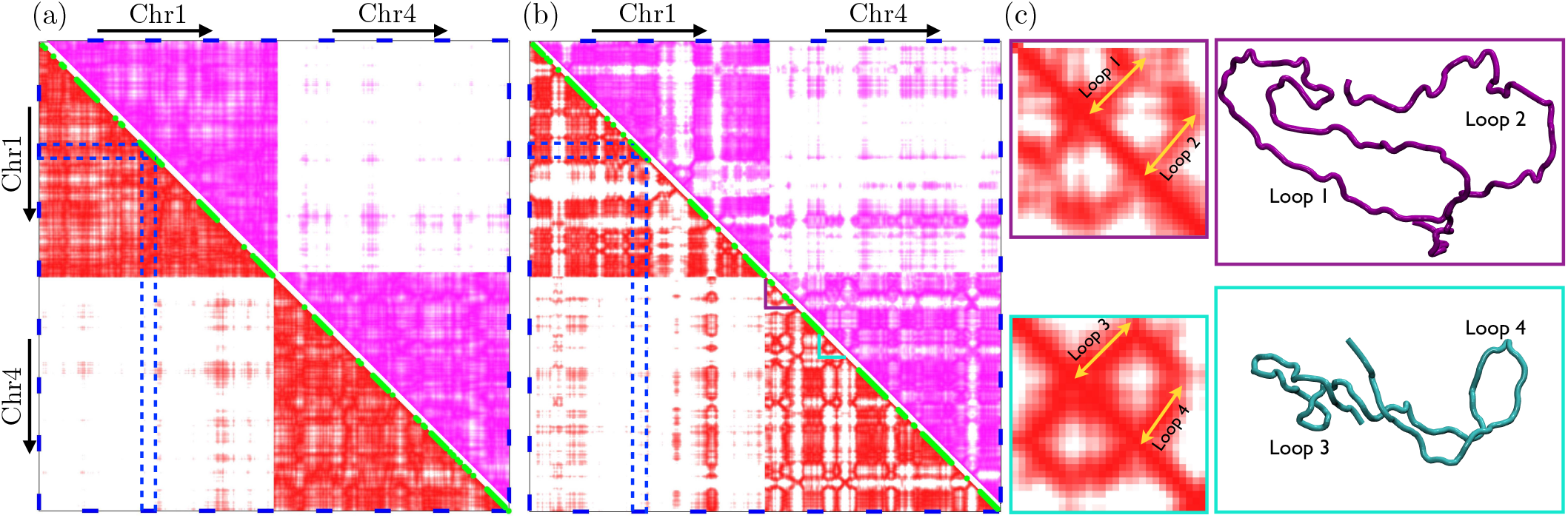
Hi-C proximity maps for (a) a passive euchromatin system (*σ*_0_ = 0) and (b) an active euchromatin system (extensile, *σ*_0_ = 30); see Appendix B for details of the method. The maps show chromosomes 1 and 4 that were not in direct contact in the initial condition (pink and orange chains in Fig. 2(d)). In each case, the lower diagonal (red) shows a time snapshot, whereas the upper diagonal (pink) shows an average over *t∈* [350, 700]. Blue segments along the axes indicate the HC blocks (four per chromosomes), which are the locations where crosslinks can form. Green segments along the diagonal show sections of the chain that are spatially contained inside the HCRs identified in Fig. 2(c). (c) Two examples of loops forming inside euchromatin in the extensile case and a zoom on their signature in the Hi-C map of (b), where their location is highlighted by purple and cyan triangles along the diagonal of chromosome 4 in (b). These panels use the data from the simulations in Fig. 2.

On the one hand, the fraction of the genome that is contained inside HCRs has a higher probability of interaction with itself due to its compacted nature; on the other hand, euchromatin away from HCRs tends to be unfolded and thus has little interaction, beyond a few links in distance, with itself or with the chromatin inside HCRs. This results in a checkerboard patterning of high and low interaction. The diagonal squares in our simulations are one signature of the highly crosslinked HCRs, which have high inter-region interaction frequency. The variable size of those squares in Fig. 5(b) reflects the number of sequential HC blocks that are occupants of the same HCR as well as the amount of trapped euchromatin. The off-diagonal squares arise from genomically distant HC blocks being bound together into a single HCR. We find that extensile activity also promotes inter-chromosomal interactions as a result of mixing, as evidenced by the off-diagonal blocks in the Hi-C maps. Checkerboard patterning, at a resolution of a 0.1 *−*1 Mb, is a prominent feature of experimental Hi-C maps and is characteristic of the spatial compartmentalization of two main types of chromatin: active and open euchromatin and inactive and compacted heterochromatin [51, 53–56]. Clear square patterns are absent in the passive simulation (*σ*_0_ = 0), where crosslinked heterochromatin forms throughout the nucleus and remains evenly distributed (see Fig. 2), resulting in a relatively stronger probability of interaction with neighboring euchromatin.

Another prominent feature evidenced in Fig. 5(b) is the formation of temporary loops inside euchromatin regions. In experimental Hi-C maps, the evidence of loops is also commonplace, where they are formed by active proteins known as loop-extruding factors such as condensin and cohesin [57, 58]. Here, the loops we observe have a different origin and derive from the unfolding of non-crosslinked chain segments by active hydrodynamic flows. In some cases, these loops occur in pairs as highlighted in Fig. 5(c), showing two examples of loop pairs and close-ups of their signatures in the Hi-C map of Fig. 5(b), where they appear as closed semi-circles along the diagonal. These structures bear resemblance to the star-shaped Hi-C patterns observed by Brandão *et al*. [59], where they were explained as a consequence of specific interactions between two condensin motors. Each yellow arrow emanating from the diagonal in the detailed Hi-C maps of Fig. 5(c) describes one loop, whereas the lines parallel to the diagonal and bridging the two arrows correspond to the adjoining bases of the two loops. The size and intensity of each semi-circle depends on the length of the loops and their compactness. We note that these patterns do not persist on long time scales and indeed disappear in the time-averaged Hi-C maps due the dynamic nature of the loops, whose fate is governed by the nucleoplasmic flows.

In summary, the proximity maps of Fig. 5 recapitulate many qualitative features observed in experimental Hi-C maps, from checkerboard patterns resulting from compaction of HCRs, to the formation of transient loops inside euchromatin due to the stretching and unfolding of the chromatin by the nucleoplasmic flows. ATP-powered extensile dipolar activity is found to play a key role in setting these genomic features, with consequences for gene interactions as well as the regulation of gene expression that remain to be explored.

## IV. DISCUSSION

Our simulations reveal the key role of ATP-dependent active processes and long-ranged hydrodynamic interactions in the spatial segregation of heterochromatin and euchromatin in the cell nucleus. By modeling chromatin as an active Zimm polymer, consisting of euchromatin blocks decorated with stochastic force dipoles and passive heterochromatin blocks subject to inter- and intra-chain crosslinks, we find that large-scale chromatin organization is profoundly affected by extensile euchromatic activity. This activity enhances both spatial segregation and compaction of heterochromatin, leading to formation of dense heterochromatin regions (HCRs). The segregation originates from nucleoplasmic flows, induced by active euchromatin, which lead to an increased mixing between chromosomes and, hence, intensified heterochromatin cross-linking within and between chromosomes. Nucleoplasmic flows display long-ranged coherence in systems with extensile dipoles, due to their spontaneous alignment in their own self-induced flow fields, which gives rise to the emergence of nematic alignment and unfolding of the polymer chain within euchromatin regions [36].

Our model for active stresses – stochastic force dipoles – is built on the premise that ATP-powered nuclear enzymes exert microscopic forces locally on chromatin segments that are transmitted to the nucleoplasm via viscous drag, that is, the force dipole acts entirely upon the nucleoplasm. More detailed microscopic modeling may suggest that forces are also transduced onto the chromatin strand directly. In that case, recent work in modeling cytoskeletal assemblies suggests that dense cross-linking can also yield long-range transmission of stress and material motion [60, 61]. The nuclear active events occur on the scale of individual genes in the human genome, the scale of a single bead in our model. Such events could be quite complex and involve active processes such as: RNA polymerase II translocation along DNA accompanied by the steric repulsion due to mRNA polymerization or local chromatin reorganization by chromatin remodellers and loop extruders such as cohesin and condensin [6, 56].

Despite this richness and complexity, the stress signature of each of such active events can be formally coarsegrained from first principles as driving a dipolar nucleoplasmic flow. This flow is the leading-order contribution in the multipole expansion conserving local momentum of the flow field generated by the active event at its chromatin chain segment. Presently, the type of dipole (i.e., extensile or contractile), its magnitude, duration and orientation with respect to the chromatin chain is unknown for different types of active events. Yet, the emergent properties of active polymers closely depend on the coupling of these active events to their conformational degrees of freedom [62, 63]. Such knowledge could be gleaned by performing microscopic molecular-scale simulations of protein-chromatin interactions and systematically coarse-graining the stress and flow fields they generate.

In our simulations, we observe heterochromatin to segregate in the center of the nucleus – a likely consequence of the symmetry of the initial data, where a randomly placed HC block is more likely to find and bind to another HC block in the nuclear interior as opposed to the periphery. This central heterochromatin placement is consistent with earlier equilibrium simulations of heterochromatin formation [25, 26]. Yet, the heterochromatin distribution observed in experiments is more complex, with heterochromatin regions located in the nuclear interior, but also along the nuclear envelope and surface of subnuclear bodies such as nucleoli [7]. This suggests that other interactions, such as chromatin-lamin interactions at the nuclear envelope [28] and chromatin-nucleolus interactions at the nucleolar surface [64], affect heterochromatin localization in the nucleus. Indeed, accumulation of heterochromatin near the nuclear envelope has been observed in equilibrium simulations accounting for chromatin-lamin interactions [14, 27]. Interestingly, both the nuclear envelope and the nucleolar surface were found to undergo dynamical shape fluctuations *in vivo* [65, 66], which could also affect the heterochromatin accumulation at these boundaries [67, 68]. Furthermore, in our model we observe an increased mixing of chromosome territories compared to experimental observations [51, 69]. This likely occurs due to high levels of activity as well as absence of physical tethering of chromatin to the nuclear envelope in our simulations.

Taken together, our model demonstrates that heterochromatin segregation is driven by an intricate interplay between the local variations in chromatin’s crosslinking and activity. Our results highlight the key role of the large-scale mechanics of chromatin and nucleoplasmic fluid, which interact via active and passive stress fields distributed across the entire nucleus. The heterogeneity of chromatin organization across the nucleus suggests a complex, spatially dependent rheological behavior, with crosslinked heterochromatin behaving like an elastic porous solid and unconstrained active euchromatin behaving like an active fluid with local nematic order [7, 49]. These two segregated materials interact via mechanical stress at their interfaces as well as viscous stresses due to nucleoplasmic flows, which permeate both phases. This picture is in good agreement with recent *in-vivo* experiments by Eshghi *et al*. [70], which used a new methodology of noninvasive microrheology that employs intrinsic chromatin dynamics to map spatially resolved rheological behavior [70, 71], and revealed that fluid-like and solid-like phases, corresponding to euchromatin and heterochromatin, respectively, coexist in differentiated cell nuclei. Modeling of these two phases revealed that euchromatin behaves as a Maxwell fluid, while heterochromatin can be described by a Kelvin solid [70]. A numerical characterization of effective rheological properties in simulations could elucidate the micromechanics underlying these measurements, as well as provide a basis for developing self-consistent continuum models of nuclear mechanics that could be used for mathematical analysis as well as mean-field simulations [72].

The discussion above begs the question as to whether there is a functional role for the organized euchromatic flows generated by dipolar activity. For example, they could facilitate the distribution of the transcription machinery in the cell nucleus by advective, in addition to diffusive, transport. Revealing biological origins of the extensile activity may provide fundamental insights into biophysical mechanism(s) behind dynamical genomic interactions and their role in gene regulation. To this end, illuminating the dynamics of the genome’s self-organization holds great promise, assisted by a close interplay between experimental and theoretical approaches across different length scales.

## ACKNOWLEDGMENTS

The authors thank Robert Blackwell for assistance with code development, and Adam Lamson and Alex Rautu for useful conversations. The authors gratefully acknowledge funding from National Science Foundation Grants CMMI-1762506 (A.Z. and M.S.), CMMI-1762566 (D.S.), DMR-2004469 (M.S.) and CAREER PHY-1554880 (A.Z.). Simulations were partially performed using the Extreme Science and Engineering Discovery Environment (XSEDE) through allocation MCB200010 (D.S.).

## Appendix A: Details of the computational model

### 1. Chain mechanics: deterministic forces

The spring forces 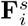 in Eq. (2) are modeled using the FENE spring law [41], which gives linear Hookean behavior at short extensions but prevents stretching beyond a maximum extension *n*_0_. Within a given chain, the net spring force on bead *i* is

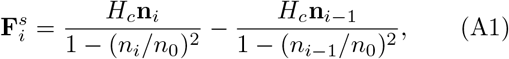

where **n**_*i*_ = **x**_*i*+1_ *−* **x**_*i*_ is the connector from bead *i* to *i* + 1 and *H*_*c*_ is the entropic spring constant. Crosslink forces 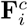 are captured using the same force law, but with a stiffer spring constant of 10*H*_*c*_.

Excluded volume forces 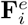 prevent overlap of distinct sections of the polymer and are captured using a soft repulsive potential:

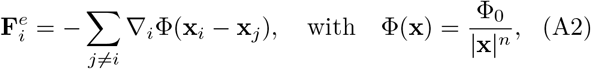

with *n* = 3. The choice of the parameter Φ_0_ is ad hoc and chosen to ensure that the chain does not cross itself. The potential is truncated whenever |**x**_*i*_ *−* **x**_*j*_| *> 𝓁*_*s*_, where *𝓁*_*s*_ is the equilibrium spring length, and three additional equally-spaced repulsive nodes are placed along each link in addition to the beads when calculating these interactions. A cell list algorithm is employed for efficient calculation of these forces with *O*(*N*) complexity [73].

### 2. Hydrodynamic interactions

The calculation of the nucleoplasmic flow field **u** appearing in the Langevin equation (1) involves solving the Stokes equation (4) subject to the no-slip condition on the nuclear envelope and evaluating the solution at the location of each bead. Due to the linearity of the Stokes equations, that solution can be written in terms of a Green’s function **G** as

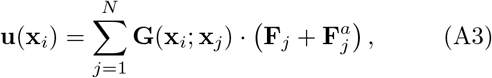

where the forces on the right-hand side are known. While analytical expressions for **G** exist in spherical domains [36, 74], a numerical solution is required in the spheroidal domains considered here. Rather than evaluating **G** directly, we instead decompose the velocity of Eq. (A3) into two parts [75]: **u** = **u**^*s*^ + **u**^*c*^. The first contribution is defined as the velocity induced by the distribution of point forces in free space,

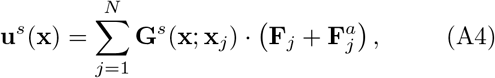

expressed in terms of the Oseen tensor or Stokeslet

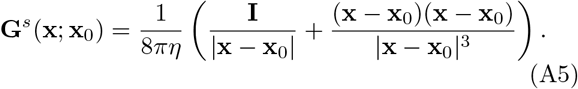

The second contribution **u**^*c*^ is a correction velocity calculated to satisfy the correct boundary condition. Note that **u**^*c*^ satisfies the homogeneous Stokes equations with boundary condition **u**^*c*^(**x**) = *−***u**^*s*^(**x**) for **x** *∈ S*. It can be calculated as a single-layer boundary integral equation for Stokes flow [76]:

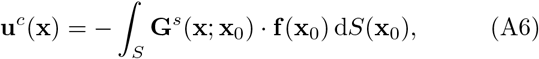

where **f** (**x**_0_) is an unknown traction distribution on the surface of the domain boundary. Evaluating Eq. (A6) on the surface *S* and using the boundary condition **u**^*c*^(**x**) = *−* **u**^*s*^(**x**) yields an integral equation for the traction field **f** that can be solved numerically. Once **f** is known on the surface for a given point force distribution, the velocity correction **u**^*c*^ is obtained by evaluating Eq. (A6) at the locations **x**_*i*_ of the beads, and can be added to the free-space velocity **u**^*s*^ of Eq. (A4) to provide the desired velocity (A3). The boundary integral equation (A6) is evaluated by quadrature after discretization of the surface into a mesh of 6-node triangular elements [76]. The algorithm was tested by comparison to the analytical Green’s function inside a spherical domain [74]. Simulations shown were performed with 5120 elements, with a numerical error of less than 2%. Finally, the calculation of both **u**^*s*^ and **u**^*c*^ is accelerated using a kernel-free fast multipole algorithm [42, 77, 78], resulting on an overall *O*(*N*) complexity with respect to the total number of beads in the system.

### 3. Thermal fluctuations

The Brownian displacements ***ξ***_*i*_(*t*) in Eq. (1) are calculated to satisfy the fluctuation-dissipation theorem, which specifies their mean and variance as

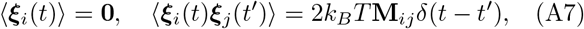

where **M**_*ij*_ denotes the grand mobility tensor that captures viscous resistance on the chain as well as long-ranged hydrodynamic interactions:

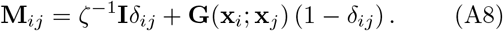

Equation (A10) is satisfied by calculating the Brownian displacements as

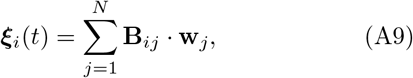

where **w**_*j*_ is an uncorrelated Gaussian white noise with zero mean and unit variance, and the tensor **B**_*ij*_ is related to the grand mobility tensor as

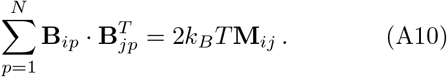

In general, solving for **B**_*ij*_ from Eq. (A10) involves either a costly Cholesky decomposition, an iterative scheme such as the Lanczos method [79], or a numerical approximation [80]. In the present work, we make the local approximation **M**_*ij*_ *≈ ζ*^*−*1^**I** *δ*_*ij*_ when calculating Brownian fluctuations, and under this approximation the right-hand side in Eq. (A10) becomes 2*D*_*b*_**I***δ*_*ij*_, where *D*_*b*_ = *k*_*B*_*T/ζ* is the Brownian diffusivity of one bead in isolation. Numerical comparison between the full solution of Eq. (A10) using the Lanczos method and the local approximation were carried out in small systems and showed negligible differences in the relevant statistical quantities.

### 4. Scalings and parameters

In all the results of Sec. III, we scale lengths with the equilibrium length *𝓁*_*s*_ of a FENE spring, forces by the corresponding spring force *F*_*s*_, and times by the characteristic time *t*_*s*_ for one isolated bead to diffuse a distance of *𝓁*_*s*_:

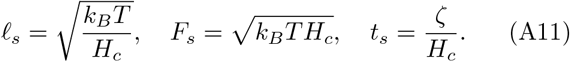

Note that *t*_*s*_ is also the spring relaxation time. Upon scaling of the system of equations, the dimensionless parameters governing the dynamics of the system are the dimensionless rate constants for active and crosslink forces, the dimensionless hydrodynamic radius *a*_*h*_*/𝓁*_*s*_, the dimensionless maximum spring extension *n*_0_*/𝓁*_*s*_, and the dimensionless active dipole strength

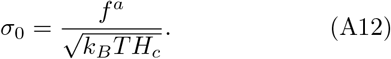

All simulations shown are for *M*_*c*_ = 23 chains of *N*_*b*_ = 1305 beads (total of 30,015 beads), with four alternating blocks of euchromatin (945 beads) and heterochromatin (360 beads) per chain, for a fraction *α*_*c*_ *≈* 0.28 of heterochromatin. The dimensionless effective radius of the nucleus is *R*_*s*_ = 28, and its eccentricity is *e* = 0.36. The hydrodynamic bead radius is *a*_*h*_*/𝓁*_*s*_ = 0.1, the maximum spring extension is *n*_0_*/𝓁*_*s*_ = 2.5, and the various dimensionless rate constants are set to 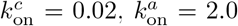, and 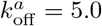. The corresponding fraction of active links along EC blocks is *p*_*a*_ = 0.285.

## Appendix B: Hi-C map calculation

The algorithm used to calculate the Hi-C proximity maps of Fig. 5 follows [51, 81] and is based on the spatial distance between pairs of genomic loci along the chromatin chains. Specifically, entries in the matrix are evaluated based on a Gaussian kernel, such that

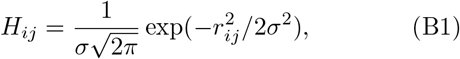

where *r*_*ij*_ = | **x**_*i*_ *−* **x**_*j*_ | is the distance between beads *i* and *j*, and where we choose a standard deviation of *σ* = 6 for the Gaussian. In practice, successive beads along the chains are binned into groups of 5, and the entries *H*_*ij*_ are averaged over each bin to generate a matrix of linear size *N*_*b*_*/*5 = 261 per chromosome. This matrix is then shown as a color plot in Fig. 5(a,b), using a linear color map from white to red (or pink) over the range of values of the matrix entries. Green dots on the diagonal in Fig. 5(a,b) highlight sections of the chromosomes that are spatially located inside HCRs, where a dot is added to the Hi-C map if at least 2 beads out of 5 inside a given bin fall inside an HCR.

## References

[1] K. E. Van Holde, Chromatin (Springer Science & Business Media, 2012).

[2] B. Alberts, A. Johnson, J. Lewis, D. Morgan, M. Raff, K. Roberts, and P. Walter, Molecular Biology of the Cell (Garland Science, 2014).

[3] J. H. Gibcus and J. Dekker, “The hierarchy of the 3D genome,” Mol. Cell 49, 773–782 (2013).

[4] B. Bonev and G. Cavalli, “Organization and function of the 3D genome,” Nat. Rev. Genet. 17, 661–678 (2016).

[5] M. R. Hübner and D. L. Spector, “Chromatin dynamics,” Annu. Rev. Biophys. 39, 471–489 (2010).

[6] A. Zidovska, “The self-stirred genome: large-scale chromatin dynamics, its biophysical origins and implications,” Curr. Op. Gen. Dev. 61, 83–90 (2020).

[7] A. Zidovska, “The rich inner life of the cell nucleus: dynamic organization, active flows, and emergent rheology,” Biophys. Rev. 12, 1093–1106 (2020).

[8] I. Solovei, K. Thanisch, and Y. Feodorova, “How to rule the nucleus: divide et impera,” Curr. Opin. Cell Biol. 40, 47–59 (2016).

[9] J. Dekker, M. A. Marti-Renom, and L. A. Mirny, “Exploring the three-dimensional organization of genomes: interpreting chromatin interaction data,” Nat. Rev. Genet. 14, 390–403 (2013).

[10] W. A. Bickmore and B. van Steensel, “Genome architecture: domain organization of interphase chromosomes,” Cell 152, 1270–1284 (2013).

[11] J. Xu, H. Ma, J. Jin, S. Uttam, R. Fu, Y. Huang, and Y. Liu, “Super-resolution imaging of higher-order chromatin structures at different epigenomic states in single mammalian cells,” Cell Rep. 24, 873–882 (2018).

[12] D. Amiad-Pavlov, D. Lorber, G. Bajpai, A. Reuveny, F. Roncato, R. Alon, S. Safran, and T. Volk, “Live imaging of chromatin distribution reveals novel principles of nuclear architecture and chromatin compartmentalization,” Sci. Adv. 7, eabf6251 (2021).

[13] I. Solovei, M. Kreysing, C. Lanctôt, S. Kösem, L. Peichl, T. Cremer, J. Guck, and B. Joffe, “Nuclear architecture of rod photoreceptor cells adapts to vision in mammalian evolution,” Cell 137, 356–368 (2009).

[14] Y. Feodorova, M. Falk, L. A. Mirny, and I. Solovei, “Viewing nuclear architecture through the eyes of nocturnal mammals,” Trends Cell Biol. 30, 276–289 (2020).

[15] T. Cheutin, A. J. McNairn, T. Jenuwein, D. M. Gilbert, P. B. Singh, and T. Misteli, “Maintenance of stable heterochromatin domains by dynamic HP1 binding,” Science 299, 721–725 (2003).

[16] C. Maison and G. Almouzni, “HP1 and the dynamics of heterochromatin maintenance,” Nat. Rev. Mol. Cell Biol. 5, 296–305 (2004).

[17] A. Janssen, S. U. Colmenares, and G. H. Karpen, “Heterochromatin: guardian of the genome,” Annu. Rev. Cell Dev. Bio. 34, 265–288 (2018).

[18] R. C. Allshire and H. D. Madhani, “Ten principles of heterochromatin formation and function,” Nat. Rev. Mol. Cell Biol. 19, 229 (2018).

[19] F. Zenk, Y. Zhan, P. Kos, E. Löser, N. Atinbayeva, M. Schächtle, G. Tiana, L. Giorgetti, and N. Iovino, “HP1 drives de novo 3D genome reorganization in early Drosophila embryos,” Nature 593, 289–293 (2021).

[20] D. Canzio, E. Y. Chang, S. Shankar, K. M. Kuchenbecker, M. D. Simon, H. D. Madhani, G. J. Narlikar, and B. Al-Sady, “Chromodomain-mediated oligomerization of HP1 suggests a nucleosome-bridging mechanism for heterochromatin assembly,” Mol. Cell 41, 67–81 (2011).

[21] P. J. Verschure, I. Van Der Kraan, W. De Leeuw, J. Van Der Vlag, A. E. Carpenter, A. S. Belmont, and R. Van Driel, “In vivo HP1 targeting causes largescale chromatin condensation and enhanced histone lysine methylation,” Mol. Cell. Biol. 25, 4552–4564 (2005).

[22] S. Machida, Y. Takizawa, M. Ishimaru, Y. Sugita, S. Sekine, J. Nakayama, M. Wolf, and H. Kurumizaka, “Structural basis of heterochromatin formation by human HP1,” Mol. Cell 69, 385–397 (2018).

[23] A. G. Larson, D. Elnatan, M. M. Keenen, M. J. Trnka, B. Johnston, A. L. Burlingame, D. A. Agard, S. Redding, and G. J. Narlikar, “Liquid droplet formation by HP1α suggests a role for phase separation in heterochromatin,” Nature 547, 236–240 (2017).

[24] A. R. Strom, A. V. Emelyanov, M. Mir, D. V. Fyodorov, X. Darzacq, and G. H. Karpen, “Phase separation drives heterochromatin domain formation,” Nature 547, 241–245 (2017).

[25] M. Falk, Y. Feodorova, N. Naumova, M. Imakaev, B. R. Lajoie, H. Leonhardt, B. Joffe, J. Dekker, G. Fudenberg Solovei, and L. A. Mirny, “Heterochromatin drives compartmentalization of inverted and conventional nuclei,” Nature 570, 395–399 (2019).

[26] Q. MacPherson, B. Beltran, and A. J. Spakowitz, “Bottom–up modeling of chromatin segregation due to epigenetic modifications,” Proc. Natl. Acad. Sci. USA 115, 12739–12744 (2018).

[27] Q. MacPherson, B. Beltran, and A. J. Spakowitz, “Chromatin compaction leads to a preference for peripheral heterochromatin,” Biophys. J. 118, 1479–1488 (2020).

[28] B. van Steensel and A. S. Belmont, “Lamina-associated domains: Links with chromosome architecture, heterochromatin, and gene repression,” Cell 169, 780–191 (2017).

[29] N. Ganai, S. Sengupta, and G. I. Menon, “Chromosome positioning from activity-based segregation,” Nucleic Acids Res. 42, 4145–4159 (2014).

[30] J. Smrek and K. Kremer, “Small activity differences drive phase separation in active-passive polymer mixtures,” Phys. Rev. Lett. 118, 098002 (2017).

[31] A. Zidovska, D. A. Weitz, and T. J. Mitchison, “Micronscale coherence in interphase chromatin dynamics,” Proc. Natl. Acad. Sci. USA 110, 15555–15560 (2013).

[32] R. Bruinsma, A. Y. Grosberg, Y. Rabin, and A. Zidovska, “Chromatin hydrodynamics,” Biophys. J. 106, 1871–1881 (2014).

[33] M. Di Pierro, D. A. Potoyan, P. G. Wolynes, and J. N. Onuchic, “Anomalous diffusion, spatial coherence, and viscoelasticity from the energy landscape of human chromosomes,” Proc. Natl. Acad. Sci. USA 115, 7753–7758 (2018).

[34] L. Liu, G. Shi, D. Thirumalai, and C. Hyeon, “Chain organization of human interphase chromosome determines the spatiotemporal dynamics of chromatin loci,” PLoS Comput. Biol. 14, e1006617 (2018).

[35] G. Shi, L. Liu, C. Hyeon, and D. Thirumalai, “Interphase human chromosome exhibits out of equilibrium glassy dynamics,” Nat. Commun. 9, 1–13 (2018).

[36] D. Saintillan, M. J. Shelley, and A. Zidovska, “Extensile motor activity drives coherent motions in a model of interphase chromatin,” Proc. Natl. Acad. Sci. USA 115, 11442–11447 (2018).

[37] A. Mahajan and D. Saintillan, “Self-induced hydrodynamic coil-stretch transition of active polymers,” Phys. Rev. E 105, 014608 (2022).

[38] D. Saintillan and M. J. Shelley, “Active suspensions and their nonlinear models,” C. R. Physique 14, 497–517 (2013).

[39] A. R. Strom, R. J. Biggs, E. J. Banigan, X. Wang Chiu, C. Herman, J. Collado, F. Yue, J. C. R. Politz, J. Tait, D. Scalzo, A. Telling, M. Groudine, C. P. Brangwynne, J. F. Marko, and A. D. Stephens, “HP1α is a chromatin crosslinker that controls nuclear and mitotic chromosome mechanics,” eLife 10, e63972 (2021).

[40] H. C. Öttinger, Stochastic Processes in Polymeric Fluids (Springer, 1996).

[41] R. B. Bird, R. C. Armstrong, and O. Hassager, Dynamics of Polymeric Liquids. Vol. 1: Fluid Mechanics (Wiley-Interscience, 1987).

[42] D. Malhotra and G. Biros, “PVFMM: A parallel kernel independent FMM for particle and volume potentials,” Commun. Comput. Phys. 18, 808–830 (2015).

[43] See Supplemental Material available at (link to be inserted by publisher) for videos of simulations corresponding to Figs. 2 and 4(d).

[44] M. V. Tamm, L. I. Nazarov, A. A. Gavrilov, and A. V. Chertovich, “Anomalous diffusion in fractal globules,” Phys. Rev. Lett. 114, 178102 (2015).

[45] D. Saintillan and M. J. Shelley, “Instabilities, pattern formation, and mixing in active suspensions,” Phys. Fluids 20, 123304 (2008).

[46] T. Gao, M. Betterton, A. Jhang, and M. Shelley, “Analytical structure, dynamics, and reduction of a kinetic model of an active fluid,” Phys. Rev. Fluids 2, 093302 (2017).

[47] A. Doostmohammadi, J. Ignés-Mullol, J. M. Yeomans, and F. Sagués, “Active nematics,” Nat. Commun. 9, 3246 (2018).

[48] Y. Hatwalne, S. Ramaswamy, M. Rao, and R. Aditi Simha, “Rheology of active-particle suspensions,” Phys. Rev. Lett. 92, 118101 (2004).

[49] D. Saintillan, “Rheology of active fluids,” Annu. Rev. Fluid Mech. 50, 563–592 (2018).

[50] J. H. Irving and J. G. Kirkwood, “The statistical mechanical theory of transport processes. IV. The equations of hydrodynamics,” J. Chem. Phys. 18, 817 (1950).

[51] E. Lieberman-Aiden, N. L. Van Berkum, L. Williams Imakaev, T. Ragoczy, A. Telling, I. Amit, B. R. Lajoie, P. J. Sabo, M. O. Dorschner, R. Sandstrom, B. Bernstein, M. A. Bender, M. Groudine, A. Gnirke, J. Stamatoyannopoulos, L. A. Mirny, E. S. Lander, and J. Dekker, “Comprehensive mapping of long-range interactions reveals folding principles of the human genome,” Science 326, 289–293 (2009).

[52] J. R. Dixon, S. Selvaraj, F. Yue, A. Kim, Y. Li, Y. Shen, M. Hu, J. S. Liu, and B. Ren, “Topological domains in mammalian genomes identified by analysis of chromatin interactions,” Nature 485, 376–380 (2012).

[53] A. Goloborodko, J. F. Marko, and L. A. Mirny, “Chromosome compaction by active loop extrusion,” Biophys. J. 110, 2162–2168 (2016).

[54] G. Fudenberg, M. Imakaev, C. Lu, A. Goloborodko, N. Abdennur, and L. A. Mirny, “Formation of chromosomal domains by loop extrusion,” Cell Rep. 15, 2038–2049 (2016).

[55] G. Fudenberg, N. Abdennur, M. Imakaev, A. Goloborodko, and L. A. Mirny, “Emerging evidence of chromosome folding by loop extrusion,” Cold Spring Harb. Symp. Quant. Biol. 82, 45–55 (2017).

[56] L. A. Mirny, M. Imakaev, and N. Abdennur, “Two major mechanisms of chromosome organization,” Curr. Opin. Cell Biol. 58, 142–152 (2019).

[57] A. L. Sanborn, S. S. P. Rao, S.-C. Huang, N. C. Durand, M. H. Huntley, A. I. Jewett, I. D. Bochkov, D. Chinnappan, A. Cutkosky, J. Li, et al., “Chromatin extrusion explains key features of loop and domain formation in wild-type and engineered genomes,” Proc. Nat. Acad. Sci. U.S.A. 112, E6456–E6465 (2015).

[58] J. Nuebler, G. Fudenberg, M. Imakaev, N. Abdennur, and L. A. Mirny, “Chromatin organization by an interplay of loop extrusion and compartmental segregation,” Proc. Natl. Acad. Sci. USA 115, E6697–E6706 (2018).

[59] H. B. Brandäo, Z. Ren, X. Karaboja, L. A. Mirny, and X. Wang, “DNA-loop extruding SMC complexes can traverse one another in vivo,” Nat. Struct. Mol. Biol. 28, 642–651 (2021).

[60] S. Fürthauer, B. Lemma, P. J. Foster, S. C. Ems-McClung, C.-H. Yu, C. E. Walczak, Z. Dogic, D. J. Needleman, and M. J. Shelley, “Self-straining of actively crosslinked microtubule networks,” Nat. Phys. 15, 1295–1300 (2019).

[61] S. Fürthauer, D. J. Needleman, and M. J. Shelley, “A design framework for actively crosslinked filament networks,” New J. Phys. 23, 013012 (2021).

[62] R. G. Winkler and G. Gompper, “The physics of active polymers and filaments,” J. Chem. Phys. 153, 040901 (2020).

[63] M. R. Shaebani, A. Wysocki, R. G. Winkler, G. Gompper, and H. Rieger, “Computational models for active matter,” Nature Rev. Phys. 2, 181–199 (2020).

[64] C. M. Caragine, S. C. Haley, and A. Zidovska, “Nucleolar dynamics and interactions with nucleoplasm in living cells,” Elife 8, e47533 (2019).

[65] F.-Y. Chu, S. C. Haley, and A. Zidovska, “On the origin of shape fluctuations of the cell nucleus,” Proc. Natl. Acad. Sci. U.S.A. 114, 10338–10343 (2017).

[66] C. M. Caragine, S. C. Haley, and A. Zidovska, “Surface fluctuations and coalescence of nucleolar droplets in the human cell nucleus,” Phys. Rev. Lett. 121, 148101 (2018).

[67] J. A. Eaton and A. Zidovska, “Structural and dynamical signatures of local DNA damage in live cells,” Biophys. J. 118, 2168–2180 (2020).

[68] M. Wang, K. Zinga, A. Zidovska, and A. Y. Grosberg, “Tethered tracer in a mixture of hot and cold brownian particles: can activity pacify fluctuations?” Soft Matter 17, 9528–9539 (2021).

[69] T. Cremer and C. Cremer, “Chromosome territories, nuclear architecture and gene regulation in mammalian cells,” Nature Rev. Gen. 2, 292–301 (2001).

[70] I. Eshghi, J. A. Eaton, and A. Zidovska, “Interphase chromatin undergoes a local sol-gel transition upon cell differentiation,” Phys. Rev. Lett. 22, 228101 (2021).

[71] C. M. Caragine, N. Kanellakopoulos, and A. Zidovska, “Mechanical stress affects dynamics and rheology of the human genome,” Soft Matter 18, 107–116 (2022).

[72] M. Theillard and D. Saintillan, “Computational meanfield modeling of confined active fluids,” J. Comp. Phys. 397, 108841 (2019).

[73] M. P. Allen and D. J. Tildesley, Computer Simulation of Liquids (Oxford University Press, 1989).

[74] C. Maul and S. Kim, “Image of a point force in a spherical container and its connection to the Lorentz reflection formula,” J. Eng. Math. 30, 119–130 (1996).

[75] J. P. Hernández-Ortiz, J. J. de Pablo, and M. D. Graham, “Fast computation of many-particle hydrodynamic and electrostatic interactions in a confined geometry,” Phys. Rev. Lett. 98, 140602 (2007).

[76] C. Pozrikidis, Boundary Integral and Singularity Methods for Linearized Viscous Flow (Cambridge University Press, 1992).

[77] L. Ying, G. Biros, and D. Zorin, “A kernel-independent adaptive fast multipole algorithm in two and three dimensions,” J. Comput. Phys. 196, 591–626 (2004).

[78] E. Nazockdast, A. Rahimian, D. Zorin, and M. Shelley, “A fast platform for simulating semi-flexible fiber suspensions applied to cell mechanics,” J. Comput. Phys. 329, 173–209 (2017).

[79] T. Ando, E. Chow, Y. Saad, and J. Skolnick, “Krylov subspace methods for computing hydrodynamic interactions in Brownian dynamics simulations,” J. Chem. Phys. 137, 064106 (2012).

[80] M. Fixman, “Construction of Langevin forces in the simulation of hydrodynamic interaction,” Macromolecules 19, 1204–1207 (1986).

[81] N. Sauerwald, S. Zhang, C. Kingsford, and I. Bahar, “Chromosomal dynamics predicted by an elastic network model explains genome-wide accessibility and long-range couplings,” Nucleic Acids Res. 45, 3663–3673 (2017).

